# mRNA-delivered neutralizing antibodies confer protection against SARS-CoV-2 variant in the lower and upper respiratory tract

**DOI:** 10.1101/2025.04.28.650951

**Authors:** Nicholas C. Hazell, Rachel A. Reyna, Awadalkareem Adam, Srinivasa Reddy Bonam, Jiani Bei, Naveen Kumar, Tina Nugyen, Jessica A. Plante, David H. Walker, Tian Wang, Kenneth S. Plante, Haitao Hu

## Abstract

Monoclonal antibodies (mAbs) have been developed as effective biological countermeasures against a range of human diseases. The high cost of antibody production and manufacturing limits its clinical application and widespread use. The mRNA-lipid nanoparticle (mRNA-LNP) is a versatile platform for development of vaccines and protein-replacement therapeutics. Since the COVID-19 pandemic, a number of neutralizing mAbs against SARS-CoV-2 have been identified with several being used clinically under emergency authorization. Herein, we report the design and generation of mRNA-LNPs expressing two SARS-CoV-2 neutralizing mAbs, 76E1 and LY1404, which respectively target the viral spike protein’s fusion peptide (FP) epitope within the S2 subunit and the receptor-binding domain (RBD) within the S1 subunit. We show a single intramuscular administration of mRNA-LNPs results in efficient LY1404 and 76E1 mAb production in mice which is sustained for 7-14 days. Further, we evaluate the protective efficacy of mRNA-LNP formulations encoding the two antibodies in mouse and hamster models challenged with different SARS-CoV-2 viral strains. The data demonstrate that a single administration of mRNA-LNP encoding the more broadly neutralizing antibody 76E1 confers significant protection against the immune-evasive SARS-CoV-2 Omicron variant BQ.1 in both the upper and lower respiratory tract of the hamsters, indicating its potential impact on limiting both viral disease and viral acquisition. Together, our study expands the potential of the mRNA-LNP platform to deliver therapeutic antibodies for rapid prevention or treatment of pathogenic infections.

## INTRODUCTION

Since the first monoclonal antibody (mAb) was granted licensure by the FDA for treating kidney transplant rejection in 1986 ^(1)^, therapeutic mAbs have begun becoming vastly manufactured as biological countermeasures for a spectrum of human afflictions, including infectious diseases ^(2)^. While methods for mAb generation have been improved considerably over the last several decades ^(3, 4)^, their production and manufacturing remains costly and laborious, which limits the widespread use and global deployment of therapeutic antibodies ^(5) (6) (7) (8)^.

Severe acute respiratory syndrome coronavirus 2 (SARS-CoV-2) has caused a major pandemic (COVID-19) by transmission via the respiratory tract ^(9–11)^. To date, a number of SARS-CoV-2 neutralizing antibodies have been identified from infected or vaccinated individuals and these antibodies target different regions of the viral spike protein with various breadths of neutralization ^(12, 13)^. LY1404, also known as bebtelovimab, is a potent neutralizing antibody targeting the receptor-binding domain (RBD) within the S1 subunit of the SARS-CoV-2 spike protein ^(14)^. It demonstrated robust neutralizing potency against early SARS-CoV-2 strains and was clinically authorized under emergency use for treating COVID-19 patients, prior to its termination due to subsequent emergences of modern variants ^(14)^. SARS-CoV-2 has constantly mutated its key epitopes in RBD, leading to the emergence of novel variants that wield an enhanced ability to evade neutralizing antibodies ^(15, 16)^. In contrast, the S2 subunit of the spike protein, compared to the S1 subunit and RBD, is genetically more conserved across human coronaviruses (HCoV), including the various SARS-CoV-2 variants ^(14)^. Multiple mAbs targeting these conserved epitopes within the S2 subunit ^(17–19)^ have also been isolated from COVID-19 convalescent patients. Compared to mAbs specific to the S1 subunit/RBD, some S2-targeting mAbs are reported to afford broader neutralization not only against SARS-CoV-2 variants, but also against other HCoVs^(14)^. One example is 76E1, a broadly neutralizing mAb that targets the fusion peptide (FP), a small cryptic epitope hidden within the S2 subunit ^(20)^. It was reported that 76E1 recognizes FP epitopes during viral infection and exhibits broad neutralization activities against immune evasive SARS-CoV-2 variants and multiple HCoVs ^(20)^.

mRNA-lipid nanoparticles (mRNA-LNP) are a versatile platform to develop new prophylactic and therapeutic countermeasures, such as vaccines, protein replacement therapy, and gene therapy against various human diseases ^(21–26)^. The two mRNA-based COVID-19 vaccines were clinically approved for use in large human populations, representing a major success in clinical application of this technology ^(27, 28)^. In addition to vaccines, mRNA-LNPs delivering therapeutic antibodies to combat infectious diseases have also been reported ^(29–33)^. Yet, it remains less clear whether mRNA-delivered antibodies targeting conserved epitopes of SARS-CoV-2 spike, e.g., the S2 FP-targeting antibody 76E1, aside from the S1/RBD-targeting antibodies, confer rapid protection against infection by highly immune evasive Omicron variants. In addition, whether systemic administration of mRNA-LNP encoding the target neutralizing antibodies can offer protection in the upper respiratory tract, thus reducing the potential risk of viral acquisition, other than in the lower respiratory system (lungs), is also unknown. These questions help delineate if mRNA-delivered neutralizing antibodies could be a promising alternative to vaccines for offering more rapid (albeit shorter) protection against contagious infections for high-risk individuals, such as travelers to endemic areas and immune-compromised populations.

In this study, we designed and generated nucleoside-modified mRNA constructs expressing the heavy chain (HC) and light chain (LC) of 76E1 and LY1404 antibodies. These mRNAs were formulated in LNPs for *in vivo* delivery in animal models. Our data show that systemic intramuscular (I.M.) administration of mRNA-LNPs encoding HC and LC of each antibody leads to efficient production of LY1404 and 76E1 *in vivo* in mice, which sustains 7-14 days in the mouse sera after single mRNA-LNP injection. We further evaluated protective efficacy of these two mRNA-delivered antibodies in mouse and hamster models that were challenged with different SARS-CoV-2 viral strains. We demonstrated that a single I.M. administration of mRNA-LNPs encoding 76E1, but not LY1404, confers significant protection against the immune-evasive SARS-CoV-2 variant BQ.1 in both lower and upper respiratory tracts of the hamsters.

## RESULTS

### Design and *in vitro* characterization of mRNAs encoding LY1404 and 76E1 antibodies

LY1404 and 76E1 are two SARS-CoV-2 neutralizing mAbs that respectively recognize epitopes within the SARS-CoV-2 spike S1 subunit (RBD) and S2 subunit (FP) (**Figure 1A**) ^(14, 20)^. To generate mRNAs expressing these mAbs, the HC and LC sequences of LY 1404 or 76E1 were codon-optimized and cloned into DNA constructs that were optimized for mRNA transcription and translation ^(34)^. Both HC and LC sequences contain a signal peptide (SP) upstream of the variable (VH and VL) and constant (CH and CL) regions to promote protein expression and secretion (**Figure 1A**). The cloned HC and LC DNA vectors were linearized, followed by mRNA synthesis using *in vitro* transcription (IVT). The mRNAs were purified through two rounds of purification, including initial precipitation by lithium chloride and the subsequent purification via a cellulose-based method to remove residual dsRNA contaminants generated during IVT ^(35)^. Gel electrophoresis confirmed the size and quality of the purified HC and LC mRNAs for both LY1404 and 76E1 mAbs (**Figure 1B**). To evaluate antibody expression, HC and LC mRNAs for either LY1404 or 76E1 mAb were co-transfected (1:1 molar ratio) into Chinese hamster ovary (CHO) cells and antibody production in culture supernatants was measured at 24 hours post-transfection (**Figure 1C**). Western blot analysis revealed detection of fully assembled LY1404 and 76E1 Abs following mRNA transfections (**Figure 1D**). Next, ELISA assays were performed on the same cell supernatants to confirm antibody expression and antigen recognition specificity. First, ELISA plates were coated with SARS-CoV-2 full-length spike protein (S), RBD, or the 15-*mer* FP peptides to examine antigen binding specificity of antibodies produced in supernatants after mRNA transfections. The data showed that LY1404, but not 76E1, effectively binds to RBD, whereas 76E1, but not LY1404, effectively binds to FP; both antibodies produced after mRNA transfection effectively bind to the full-length S protein **(Figure 1E)**. Further, LY1404 and 76E1 antibodies in supernatants were quantified using a human IgG ELISA kit (capture antibody specific for human IgG). We observed that both antibodies were readily detected at high levels following HC and LC mRNA co-transfections **^(Figure 1F)^**. These data support that the HC and LC mRNAs can be combined at a 1:1 molar ratio for efficient production of LY1404 and 76E1 mAbs in cells.

**Figure 1.**
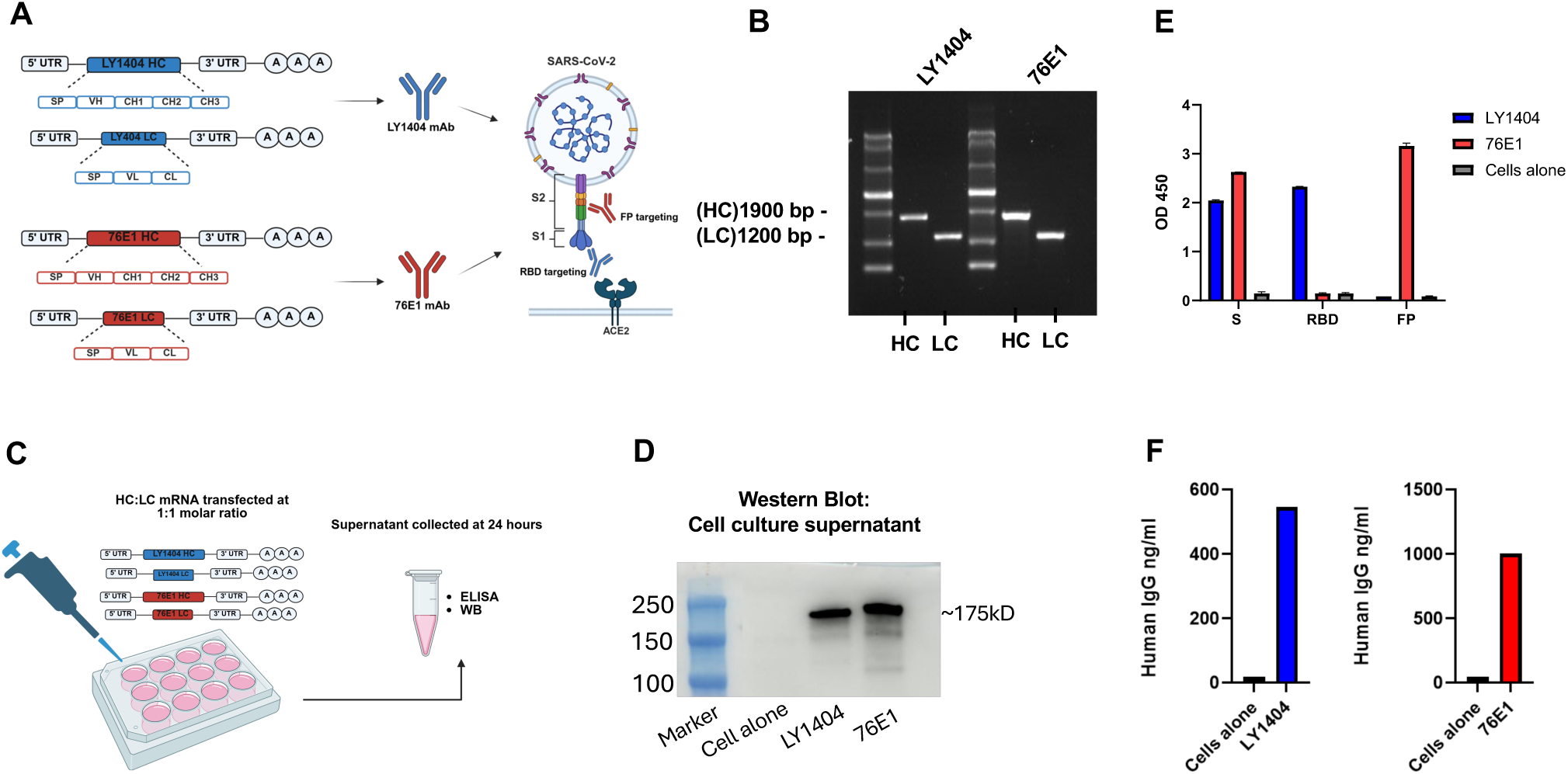
Generation and *in vitro* characterization of mRNAs encoding LY1404 and 76E1 HC and LC. **(A)** Design of mRNA constructs expressing LY1404 (blue) or 76E1 (red) heavy chain (HC) and light chain (LC). A graphical representation of the targeted regions in the SARS-CoV-2 S protein by both mAbs is also shown. In the mRNA constructs, SP: signal peptide; VH: heavy chain variable; CH: heavy chain constant; VL: light chain variable; CL: light chain constant. **(B)** Gel electrophoresis analysis of the purified HC and LC mRNAs of LY1404 and 76E1. **(C)** Schematic illustration of in vitro characterization of antibody-encoding mRNAs in cells. CHO cells were transfected with HC and LC mRNA mixture (1:1 molar ratio) for each antibody and culture supernatants were collected 24 hours after transfection. Antibody production in supernatants was measured by western blot (WB) and ELISAs. **(D)** WB analysis of antibody production in supernatants. **(E)** Determination of antibody production in culture supernatants by antigen binding ELISA. ELISA plates coated with SARS-CoV-2 full-length S protein, RBD, or FP peptides, as indicated were used for the detection of individual antibody production and antigen recognition specificity. Adjusted OD450 values were shown. **(F)** Quantification of LY1404 and 76E1 antibodies in the culture supernatant using quantitative human IgG ELISA kit.

### Antibody mRNA encapsulation into LNPs for *in vivo* delivery

We next formulated the purified HC and LC mRNAs into LNPs for *in vivo* delivery. The LNP formulation was summarized in **Figure 2A**. Individual HC or LC mRNA was diluted in citrate acid buffer as the aqueous mRNA phase, which was mixed at a 3:1 flow-rate ratio with an ethanol organic phase containing four different lipids (SM102, DSPC, Cholesterol, and DMG-PEG-2K) **(Figure 2A**). Individual HC or LC mRNA-LNPs were dialyzed and concentrated, followed by characterization of LNP size, polydispersity index (PDI), and RNA encapsulation efficiency **(Figure 2B-C)**. The data showed that all four HC and LC mRNA-LNPs manifested comparable particle size (102-110nm), PDI (0.107-0.134), as well as RNA encapsulation efficiency (94%-96%) **(Figure 2C)**. The mRNA-LNPs were employed for *in vivo* testing in animal models.

**Figure 2.**
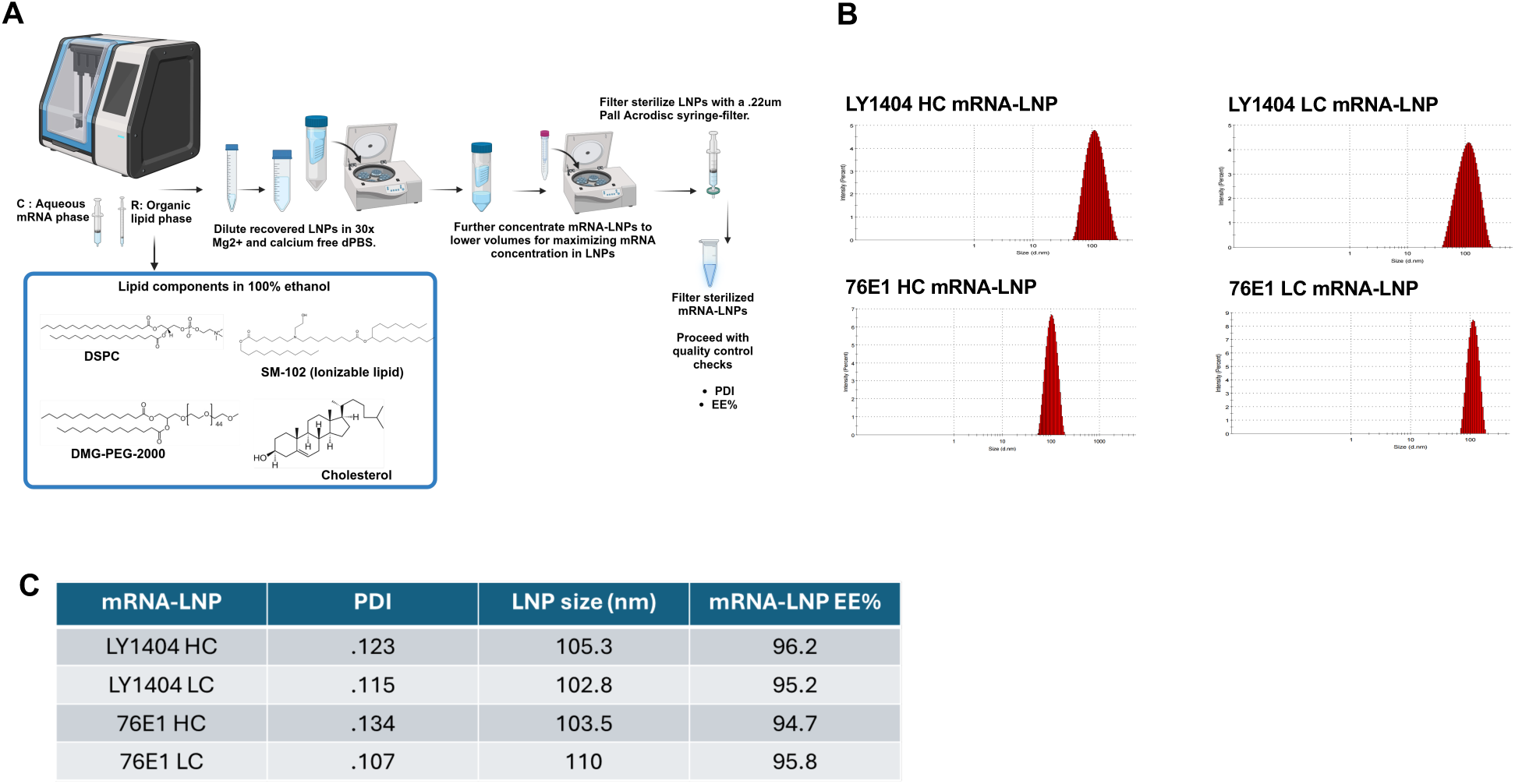
Formulation of mRNAs in LNP and mRNA-LNP characterization. **(A)** Schematic illustration of procedures for mRNA-LNP formulation and purification. Individual lipids employed for LNP formulation (SM102, DSPC, Cholesterol, and DMG-PEG2000) are shown. **(B-C)** Determination of LNP size, PDI, and mRNA encapsulation efficiency. LNP size and PDI were measured by Zetasizer. A representative graph for each mRNA-LNP PDI and size is shown (B). mRNA encapsulation efficiency and concentration in LNP were determined by Ribogreen assay. mRNA-LNP size, PDI, and RNA encapsulation efficiency were summarized in (C).

### In vivo expression and kinetics of LY1404 and 76E1 antibodies in mice

Next, to test if the HC and LC mRNA-LNPs can efficiently produce LY1404 or 76E1 antibodies *in vivo*, two groups of BALB/C mice (n=5/group) were respectively administrated with LY1404 HC and LC mRNA-LNP mixture or 76E1 HC and LC mRNA-LNP mixture. Total dose of mRNA per animal was 15µg for HC and LC mRNA together (1:1 molar ratio). mRNA-LNPs were administered intramuscularly **(Figure 3A)**. Blood was collected at 1, 3, 7, and 14 days after mRNA-LNP administration to measure target antibody production in sera **(Figure 3A)**. Blood was also collected from mice prior to the mRNA-LNP injection as a baseline measurement of antibodies. Serum antibody levels were measured by the two ELISA assays as described above **(Figure 3B-D)**. RBD binding ELISA showed that mRNA-LNP administration led to production of significant levels of LY1404 in sera 24 hours post administration, which reached peak levels at 3 days and maintained at high levels until 7 days after administration, whereas no RBD-reactive antibody was detected in the sera of mice injected with 76E1-encoding mRNA-LNPs **(Figure 3B)**. In contrast, FP binding ELISA showed that mRNA-LNP administration led to production of high levels of 76E1 antibody in sera at 24 hours after administration, which similarly reached peak levels at 3 days after administration; no FP-reactive antibody was detected in the sera of mice injected with LY1404-encoding mRNA-LNPs **(Figure 3C)**. Compared to LY1404 which remained detectable at high levels at 7 days, 76E1 antibody production appears declining to low levels at 7 days after mRNA-LNP injection with only one out of the five mice manifesting detectable antibody in sera **(Figure 3B-C)**. Further, we performed quantitative human IgG ELISA to quantify LY1404 or 76E1 antibody in the mouse sera and observed consistent results **(Figure 3D)**. Compared to the baseline sera, administration of mRNA-LNPs encoding LY1404 or 76E1 both led to detection of human IgG in mouse sera, and the kinetics of antibody levels were similar to those of RBD- or FP-binding ELISAs with maximal levels detected at 3 days after administration (500-1000ng/ml in sera) **(Figure 3D)**. By day 14 post mRNA-LNP administration, both 1404 and 76E1 antibodies became undetectable in the mouse sera **(Figure 3B-D)**. Together, these data support efficient production of antibodies with specific antigen recognition specificity *in vivo* following I.M. mRNA- LNP administration.

**Figure 3:**
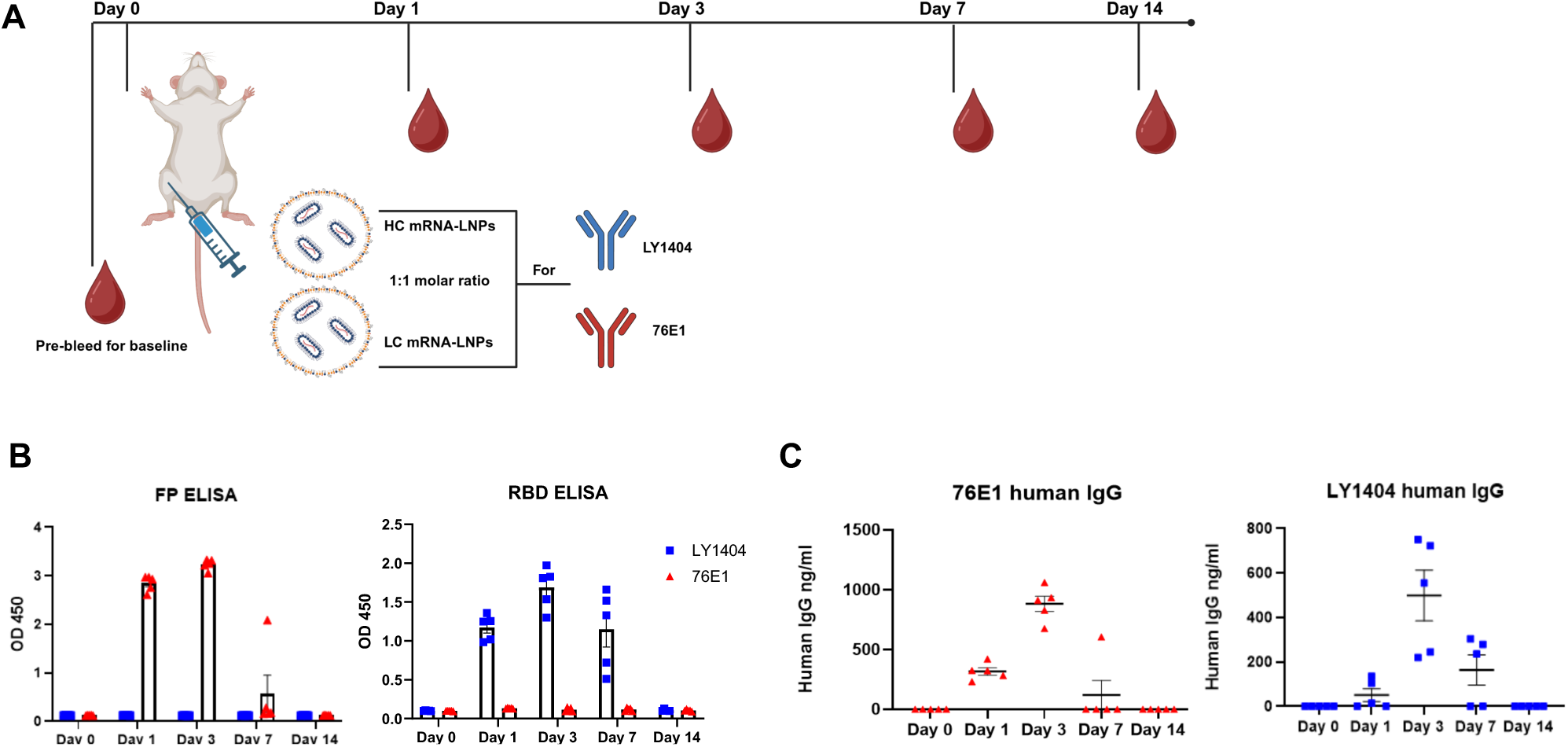
1404 and 76E1 antibody production in vivo following mRNA-LNP administration. **(A)** Mouse study design. Two groups of 6-week-old BALB/C mice (n=5/group) were intramuscularly administrated with mRNA-LNPs containing LY1404 HC and LC or 76E1 HC and LC (HC and LC mRNA at 1:1 molar ratio). Blood was collected at 1, 3, 7, and 14 days after mRNA-LNP administration to measure target antibody production in sera. Blood was also collected from mice prior to the mRNA-LNP injection to determine baseline levels. **(B)** Kinetics of serum antibody production as measured by antigen-binding ELISA. ELISA plates were coated with either FP peptides or RBD protein as indicated to determine antibody level and antigen-recognition specificity. Adjusted OD450 values for each time point were shown. **(C)** Quantification of LY1404 and 76E1 antibodies in the mouse sera using quantitative human IgG ELISA kit. Kinetics of antibody levels are shown (ng/ml).

### Protection of mRNA-encoded antibodies against mouse-adapted (MA) SARS-CoV-2 infection in mice

After confirming *in vivo* production of LY1404 and 76E1 antibodies by mRNA-LNPs, we next evaluated their protective efficacy against SARS-COV-2 infection in animal models. First, wild-type mice infected with a mouse-adapted (MA) SARS-CoV-2 CMA4 strain was employed as a less stringent challenge model. 6-week-old female BALB/C mice were intramuscularly administered with mRNA-LNP containing LY1404 or 76E1 HC and LC mRNAs (15ug at 1:1 molar ratio) as described above. Another group of mice receiving an equivalent amount of empty LNP (eLNP) was included as a negative control **(Figure 4A)**. One day after mRNA-LNP administration, all mice were intranasally challenged with CMA4 MA-SARS-CoV-2 (1 x 10^4^ pfu/mouse) **(Figure 4A)**. Two days after viral challenge, mice were euthanized for measuring infectious viral titers in the lungs **(Figure 4A)**. Compared to the mice of the control group that received eLNP, mice receiving either LY1404 or 76E1 mRNA-LNPs had a significant reduction in lung viral titers **(Figure 4B)**. Relative to eLNP, LY1404 mRNA-LNP reduced lung viral titers by 65 folds (from 755,993 pfu/g to 11,626 pfu/g; p<0.0001) and 76E1 mRNA-LNP reduced lung viral titers by 49 folds (from 755,993 pfu/g to 15,343 pfu/g; p<0.0001) (**Figure 4B**). Together, the results from MA-SARS-CoV-2 infection model provide evidence that mRNA-delivered LY1404 or 76E1 antibody inhibits SARS-CoV-2 infection in mice.

**Figure 4.**
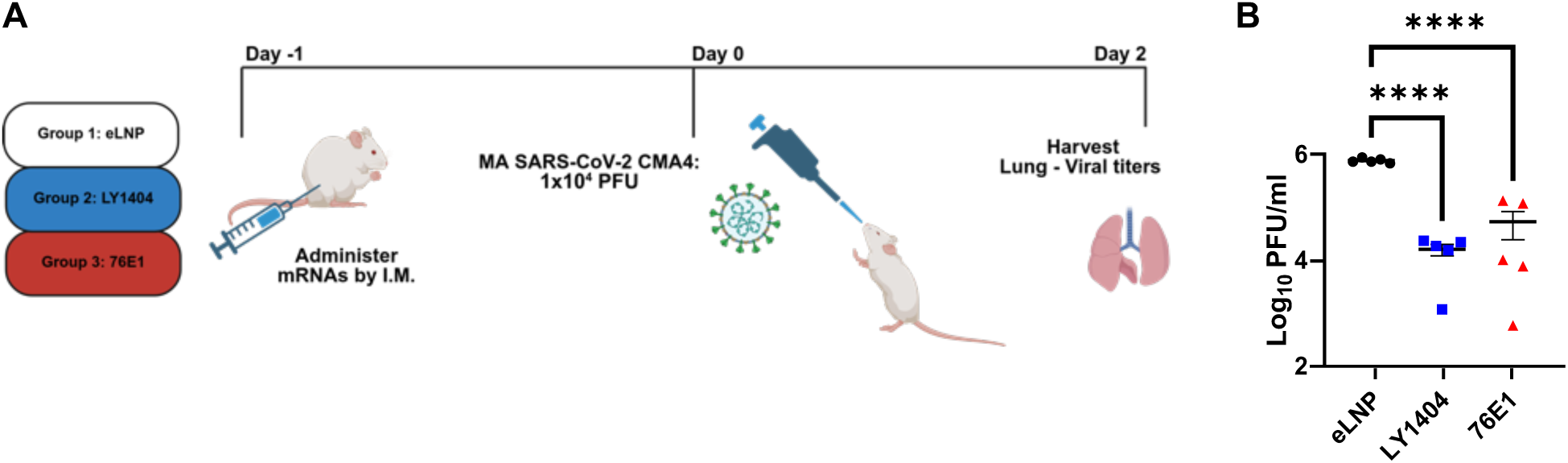
Protective efficacy of mRNA-encoded 1404 and 76E1 antibodies against SARS-CoV-2 in mice. **(A)** Mouse study design. 6-week-old female BALB/C mice (n=5/group) were intramuscularly administered with mRNA-LNP containing LY1404 or 76E1 HC and LC mRNAs (15µg at 1:1 molar ratio). Another group of mice receiving equivalent amount of eLNP was included as a negative control. One day after mRNA-LNP administration, all mice were intranasally challenged with CMA4 MA-SARS-CoV-2 (1 x 10^4^ pfu), following by terminal harvest and quantification of lung viral titers at 2 days after viral challenge. **(B)** Quantification of infectious viral titers in mouse lungs of all three groups. **** p<0.0001.

### Protection of mRNA-encoded LY1404 and 76E1 antibodies against immune evasive SARS-CoV-2 variant in hamsters

Based on the above findings, we next assessed protective efficacy of the mRNA-encoded LY1404 and 76E1 antibodies against more immune evasive SARS-CoV-2 variant. We selected Omicron BQ.1, a variant showing much increased ability to transmit and to evade antibody neutralization compared to MA-SARS-CoV-2 CMA4 and earlier SARS-CoV-2 strains ^(15, 36)^. It should be noted that RBD-targeting antibodies, including LY1404, while manifesting robust potency against early SARS-CoV-2 strains, had much reduced neutralization activity towards SARS-CoV-2 Omicron variants, like BQ.1, whereas the FP-targeting antibody 76E1 has shown broader neutralization against SARS-CoV-2 variants. To test efficacy of mRNA-encoded antibodies against BQ.1, three groups of hamsters (6-week-old; n=5 per group) were respectively administered with mRNA-LNPs encoding antibodies (LY1404, 76E1) or with eLNP as negative control. mRNA-LNPs were also given intramuscularly **(Figure 5A)** with a total mRNA dose of 15ug for each antibody (HC and LC mRNA at 1:1 molar ratio), identical to the regimens used in the above two experimental designs. One day after mRNA-LNP administration, hamsters were intranasally challenged with SARS-CoV-2 BQ.1 (2 x 10^4^ PFU/ml). Two days after viral challenge, viral titers in the hamster lungs (lower respiratory tracts; LRT) as well as in the nasal washes (NW) (upper respiratory tracts; URT) were measured to assess the impacts of mRNA-encoded antibodies on viral infections in both LRT and URT **(Figure 5A)**.

**Figure 5.**
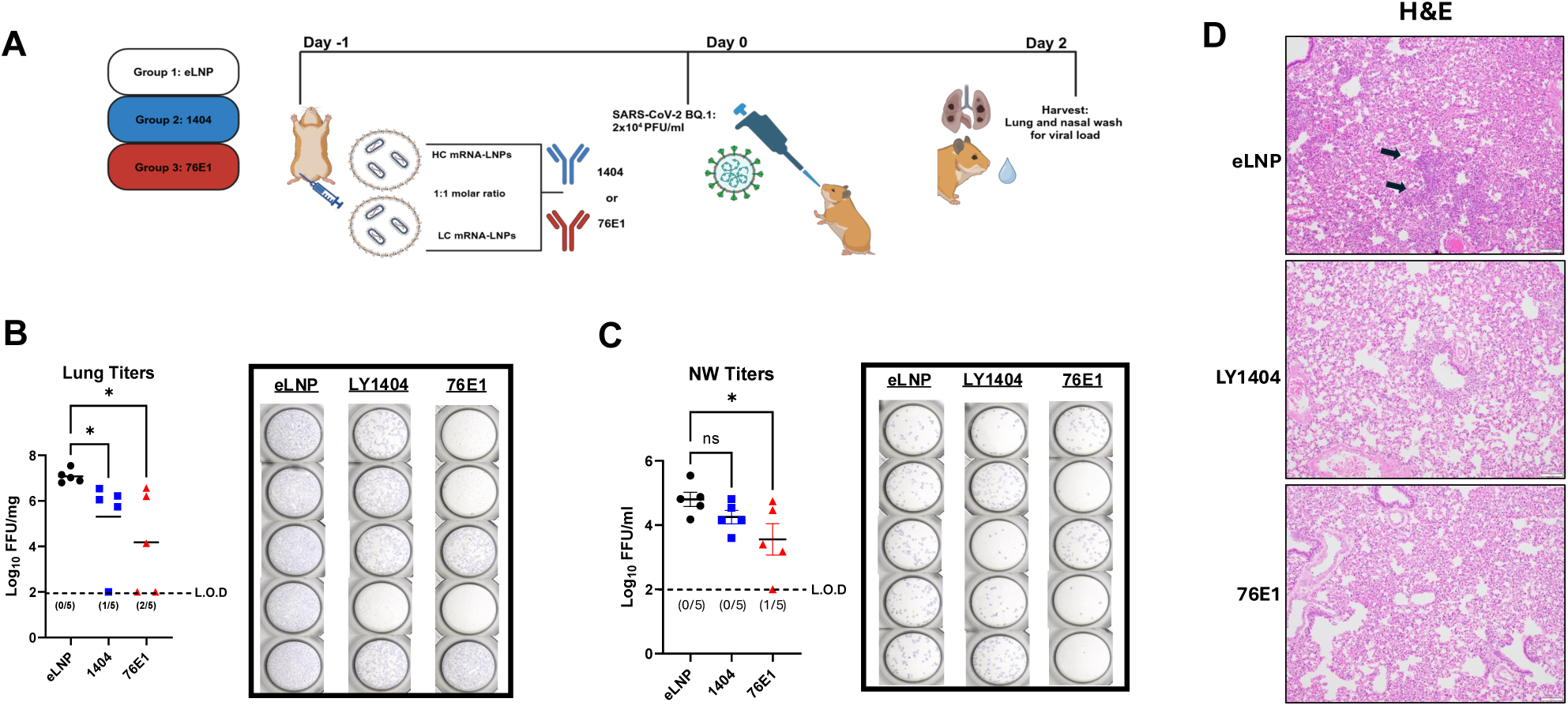
Protective efficacy of mRNA-encoded LY1404 and 76E1 antibodies against SARS-CoV-2 Omicron variant BQ.1 in hamsters. **(A)** Hamster study design. Three groups of hamsters (6-week-old; n=5 per group) were respectively administered with mRNA-LNPs encoding antibodies (LY1404, 76E1) or with eLNP as negative control. One day after mRNA-LNP administration, hamsters were intranasally challenged with SARS-CoV-2 BQ.1. Two days after viral challenge, viral titers in the hamster lungs (lower respiratory tracts; LRT) as well as in the nasal washes (NW) (upper respiratory tracts; URT) were measured. Hamster lungs were also stained for histopathological analysis. **(B-C)** Quantification of infectious viral titers in the lungs (LRT) (B) and nasal washes (NW; URT) (C) of hamsters. **(D)** Histopathological staining of hamster lungs. Representative lung H&E staining of each group is shown. * p<0.05.

We observed that both mRNA-encoded LY1404 and 76E1 antibodies induced significant protection against BQ.1 in the hamster lungs **(Figure 5B)**. Compared to the eLNP group, in which all 5 hamsters had high viral titers with geometric mean of 1.2 x 10^7^ pfu/g, administration with LY1404-encoding mRNA-LNPs led to complete viral suppression (no detectable viral titers) in one hamster and reduced viral titers in four hamsters (viral titer geometric mean: 162,690 pfu/g in LY1404 group; 86-fold reduction compared to eLNP group) (p<0.05) **(Figure 5B)**. Similarly, administration with 76E1-encoding mRNA-LNPs resulted in a robust protection against BQ.1 in hamster lungs: complete viral suppression (no detectable viral titers) in two hamsters and reduced viral titers in other three hamsters with a geometric mean of 9,487 pfu/g in this group (>1480-fold reduction compared to eLNP group) (p<0.05) (**Figure 5B**). Of note, when quantifying viral loads in the NW (URT), we observed that mRNA-encoded 76E1, but not LY1404, also significantly reduced viral titers in the URT. Compared to the eLNP group (geometric mean viral titers: 633,92 pfu/ml in NW), administration with mRNA-encoded 76E1 reduced viral titers to geometric mean of 2,274 pfu/ml in NW (28-fold reduction compared to eLNP group; p<0.05), leading complete viral suppression in the NW in one hamster **(Figure 5C)**. In contrast, administration with mRNA-encoded LY1404 only slightly reduced viral titers in the NW compared to the eLNP group with no statistical significance detected **(Figure 5C)**. Finally, hamster lung histopathological examination was performed to further evaluate protection of mRNA-encoded antibodies against viral challenge. As shown (**Figure 5D**), SARS-CoV-2 BQ.1 infection caused evident lesions in the lung of eLNP-treated control hamsters, including focal interstitial pneumonia (**Figure 5D**, top panel). Hamsters administered with either LY1404- or 76E1-encoded mRNA-LNPs were protected from lung lesions and demonstrated normal bronchial, bronchiolar, and alveolar architecture (**Figure 5D**, mid and bottom panels), which are consistent with the data on the hamster lung viral titers. Together, these data support that systemic administration of antibody-encoding mRNA-LNP (via I.M. route) confers significant protection against immune-evasive SARS-CoV-2 variant in the lungs (URT) and that the mRNA-encoded 76E1 antibody also induces significant protection against SARS-CoV-2 BQ.1 in the upper reparatory tract (URT), indicating potential impact of mRNA-encoded 76E1 Ab on viral acquisition or transmission, other than its protective effect on lung infection and pathology.

## DISCUSSION

The mRNA-LNP platform has gained much attention for its ability to deliver therapeutic proteins or antibodies as effective countermeasures against a range of diseases ^(21–24)^, in addition to its milestone success in vaccine development ^(27, 28)^. To combat viral diseases including the COVID-19 pandemic, many neutralizing antibodies with therapeutic potential were identified ^(12)^ and some were approved under emergency use. A main challenge of wide implementation of antibody therapeutics is the high cost and labor of antibody production and manufacturing. Here, we show that mRNA-LNP can effectively deliver two different SARS-CoV-2 neutralizing antibodies (LY1404 and 76E1) for expression *in vivo* and that a single dose of mRNA-LNP (15µg for combined HC and LC mRNAs) confers significant protection against SARS-CoV-2 infection in two different animal models, including use of the highly immune-evasive Omicron variant. Notably, our data also support that I.M. administration of mRNA-LNPs encoding for 76E1 significantly protects against BQ.1 not only in lungs (LRT) but also in the URT of hamsters, suggesting potential impact of the approach in limiting viral infection or transmission. Our study support that this mRNA-based approach may serve as a promising alternative to vaccines for inducing rapid, albeit short-term, protection for high-risk (e.g., travelers to endemic areas) and immune-compromised populations.

A number of human neutralizing antibodies against SARS-CoV-2 have been identified ^(12)^. Those identified earlier after the pandemic, including LY1404, mainly target the viral spike S1 subunit and RBD. However, the S1/RBD regions are more prone to mutations, leading to the emergence of many SARS-CoV-2 variants highly immune evasive to the antibody responses induced by licensed vaccines or antibody therapeutics^(15, 37)^. Indeed, LY1404 was eventually removed from the emergency authorization list due to its loss in efficacy against the immune escaping Omicron strains, namely BQ.1 and XBB ^(36)^. In contrast to LY1404, 76E1 targets the more conserved FP epitope in the S2 subunit and is considered more broadly neutralizing ^(20)^. Different groups have isolated mAbs specific to the S2 subunit and shown that they offer broader neutralization against a range of HCoVs compared to the S1-targeting mAbs ^(20, 38)^. Our study compared *in vivo* efficacy of the two representative RBD- and FP-targeting antibodies (LY1404 and 76E1) when delivered by mRNA-LNP in animal models. While both mRNA-encoded antibodies manifested comparable efficacy against the MA-SARS-CoV-2 in WT mice, only mRNA-encoded 76E1 offered significant protection against the immune-evasive BQ.1 challenge in both URT and LRT of hamsters. Thus, our results are consistent with these earlier findings and support that inclusion of S2 (or FP)- targeting mAbs should be pursued in future vaccine and therapeutic strategies to offer broader protection against viral variants and different HCoVs.

LNP delivery of mRNA payload is critical to the development of mRNA vaccines and therapeutics ^(39–41)^. In the current study, we utilized the conventional liver-oriented LNP (SM102) and performed systemic administration (intramuscular) of antibody-encoding mRNA-LNPs. The data show that I.M. injection leads to rapid detection of antibodies in the circulation within 24 hours post injection.

Consistently, we observed a significant protection against BQ.1 infection in the lungs/lower respiratory tracts of hamsters after systemic administration of 76E1 mRNA-LNPs. This could attribute to the presence of 76E1 antibodies in circulation and/or lungs following I.M. injection. Of note, we also observed significant protection against BQ.1 in the hamsters URT by 76E1 mRNA-LNPs following I.M. injection. This is of interest as SARS-CoV-2 variants like BQ.1 have wielded a stronger ability to spread by localizing more in the upper airways ^(42)^. Our data indicates the potential impact of this approach on reducing risk of viral acquisition and/or transmission in the URT. The mechanism of URT protection by systemic (I.M.) mRNA-LNP administration remains less clear in the current study and warrants further investigations in future. In addition, growing efforts have been pursued in discovering novel LNPs for organ- and site-specific mRNA delivery ^(43)^, including mucosal/intranasal delivery ^(44, 45)^ and lung delivery ^(46, 47)^. In future, it would be interesting to investigate site-specific delivery of antibody-encoding mRNAs to lungs or to nasal mucosa using these novel LNPs and compare their protective efficacy against viral challenge with the systemically administered LNPs.

Our study examined kinetics of circulating antibody production in mice following a single dose of mRNA-LNP and revealed that both LY1404 and 76E1 were rapidly detectable in the sera within 24 hours post injection. The serum antibodies peaked around 3 days and gradually diminished over time to undetectable levels by 14 days post administration. This serum antibody kinetics is consistent with a previous report using similar mRNA-LNP technology to deliver VRCO1, an HIV- 1 neutralizing antibody^(33) (33)^. The modest durability of antibody production in our animal models could be attributed to multiple factors, including the transient expression nature of mRNA and possible immune suppression of human IgG antibodies expressed in the mouse host. One way to prolong antibody expression is repeated mRNA-LNP administration ^(33)^. Alternatively, compared to mRNA, other RNA technologies such as circular RNA ^(48–50)^ and self-amplifying RNA ^(51, 52)^ may provide some advantages in delivering therapeutic proteins or antibodies in terms of RNA stability and duration of target protein expression. These technologies should be explored in future for expressing therapeutic proteins or antibodies to enhance the duration of protein expression as compared to the mRNA platform.

Limitations of the current study are noted. First, it would be ideal to include protein format of the two antibodies (LY1404 and 76E1) as controls in the animal experiments to better understand dose-sparing effect of mRNA administration. Based on a previous study testing mRNA-encoded antibodies in rodents ^(33)^ and the clinical dose of LY1404 in humans under emergency use authorization ^(14)^, we speculate hundreds of µg protein antibody will likely be needed to achieve similar protection in hamsters, as compared to the 15µg mRNA dose in our study. Second, the current study focused on the prophylactic model where mRNA-encoded antibodies were administered prior to viral challenge. Efficacy of this approach to treat animals after viral infection and duration of protection remain less clear and should be further explored. In summary, our study presents a proof-of-concept that systemic administration of mRNA-LNP encoding human neutralizing antibodies confers protection in both lower and upper respiratory tracts against immune-evasive SARS-CoV-2 variant in animal models, supporting potential utility of this approach as an alternative to vaccines to offer rapid (short-term) protection for high-risk and immune-compromised populations. The study also has implications for the development of RNA- based protein replacement or antibody therapeutics for human diseases beyond infections.

## METHODS

### mRNA synthesis and purification

mRNA synthesis and purification were performed using methods reported in our previous studies with modifications ^(34)^. In brief, the HC and LC gene sequences of LY1404 and 76E1 were codon-optimized, synthesized and cloned into a DNA vector for mRNA synthesis. Nucleoside-modified mRNAs were generated from linearized DNA vector using the HiScribe CleanCap mRNA IVT kits (NEB) along with replacing uridine with N1-methyl-psuedoUridine. mRNA was precipitated from the IVT reaction using lithium chloride, followed by cellulose-based purification as previously reported ^(35)^ to remove residual dsRNA. mRNAs were analyzed by gel electrophoresis and their concentrations were determined, followed by storage at -20C for further characterization and LNP formulation.

### ELISA and WB measurement of LY1404 and 76E1 antibody expression by cells

Approximately 2x10^5^ CHO cells were seeded in 12 well plates one day prior to mRNA transfection. HC and LC mRNA for either LY1404 or 76E1 were transfected into CHO cells at a 1:1 molar ratio using Lipofectamine MessengerMAX transfection kit (Thermo Fisher). The cell culture supernatant was collected at 24 hours post mRNA transfection to measure antibody production by ELISA. ELISA assays were conducted with binding antigens or human IgG capture antibody, using a protocol as previously reported ^(34)^. Western blot (WB) was also conducted to confirm antibody expression. In brief, supernatant samples were mixed 1:1 with Laemmli buffer followed by boiling at 95°C for 10 minutes. Full LY1404 and 76E1 IgG expression was determined using an anti-human IgG lambda LC antibody (1:3000) (Abcam). Anti-lamda LC antibodies were incubated with WB membranes in wash buffer with 5% milk overnight at 4°C. WB membranes were then washed and incubated with secondary HRP-conjugated antibody (1:3000) for 1 hour at room temperature. Membranes were washed and visualized by using enhanced chemiluminescence Western blotting substrate (Thermo Fisher Scientific).

### mRNA LNP encapsulation into LNPs

mRNAs were diluted in citrate buffer (pH 4; 20mM) to a concentration of 0.2µg/µl represented as the mRNA aqueous phase. The lipid organic phase contained ionizable lipid (SM-102), DSPC, Cholesterol, and DMG-PEG-2000 dissolved in ethanol at a molar ratio of 50:10:18.5:1.5 (vendors of individual lipids). The mRNA aqueous phase and lipid organic phase were mixed together using a NanoAssemblr microfluidic machine (GenVoy) at a 3:1 flow-rate ratio and 12ml/min total flow rate. Assembled mRNA-LNPs were dialyzed with 30X (volume ratio) dPBS containing 10% sucrose, followed by concentration using Amicon ultra-spin filters (10kDa MWCO) followed by syringe-filter sterilization using 0.22µm Pall filters. Polydispersity index (PDI) and size of the nanoparticles were measured by a Malvern dynamic light scatter (DLS) zeta-sizer. Final mRNA concentration and total encapsulation efficiency within formulated LNPs were measured by a Quant-it^tm^ RiboGreen reagent and RNA assay kit (ThermoFisher). Filter sterilized mRNA-LNPs were stored at -80°C after characterization prior to use.

### Antibody expression in mice following 15µg mRNA-LNP intramuscular administration

Antibody expression and their kinetics were tested in 6-week-old female BALB/c mice (the Jackson Laboratory; strain no. 000651). Mice were intramuscularly administered a single 15µg dose of mRNA-LNPs containing the HC and LC mRNAs encoding either LY1404 or 76E1 mAbs. Blood/sera were collected from mice prior to mRNA-LNP injection (as a baseline control), as well as 1, 3, 7, 14 days after for monitoring. Serum antibodies were measured by binding and quantitative ELISAs as detailed below.

### Binding IgG ELISA for confirming binding specificity

For confirming both antibodies recognition of the same target and their respective targets, cell culture supernatant and harvested animal serum were used in binding IgG ELISA assays. For LY1404, ELISA plates were coated overnight at 4°C with dPBS containing 1ug/ml of recombinant RBD (Sino-biological). The next day, plates were washed 2 times with PBS containing 1.5% tween followed by blocking for 1 hour at 37°C with dPBS with 8% FBS. Plates were washed two times and coated with samples supernatants or serum and incubated again at 37°C for one hour. Plates were washed five times then coated with anti-human IgG antibody (AbCam) at a 1:3000 dilution in wash buffer for 1 hour 37°C. After final wash, plates were developed using TMB 1-Component Peroxidase Substrate (ThermoFisher Scientific), followed by termination of reaction using the TMB stop solution (ThermoFisher Scientific). Plates were read at 450 nm wavelength within 15 minutes by using a Microplate Reader (BioTek).

### Detecting 76E1 specificity by using FP peptide binding ELISA assay

Considering the cryptic nature of FP, we wanted to ensure our 76E1 mRNA delivered antibody recognizes its appropriate target in specific FP epitopes. The amino acid JPT peptide-synthesizing company generated 15 *mer* overlapping peptides spanning (816- 835 aas) of the spike proteins S2 subunit. These epitopes have previously been considered the *bone-fide* FP. The FP peptides were biotinylated for use with streptavidin ELISA plates supplies from Invitrogen (Cat:15125) at a 1mM concentration. ELISA assays were performed the same day as plating. Briefly, FP peptides were diluted in peptide coating buffer at a 1mM/ml concentration and incubated for 1hour at room temperature. Plates were then washed five times and blocked with dPBS at a 10mg/ml of BSA at room temperature for 30 minutes. Blocking fluid was then removed off without washing and wand allowed to dry. Supernatant or serum samples were then added to the plate diluted in blocking buffer at a 1:20 for both sample types and incubated at 37°C for 1 hour. ELISA plates were then washed and incubated with anti-human antibody 1:3000 dilution for 1 hour at 37°C. After the final wash, developer was added and OD-450 values read on a plate-reader.

### Human IgG ELISA for quantifying mAb expression

To quantify antibody concentration and ensure it is humanized, we used a human IgG capture ELISA with a human IgG standard. ELISA plates were coated with a capture human IgG antibody in dPBS at 1:1000 and incubated 4°C over-night. The next day, ELISA plates were washed 6 times followed by blocking with dPBS containing 5% milk for 1hour at 37°C. After 1hour, blocking is dumped, no wash, with plates allowed to dry prior to adding samples. Samples were diluted 1:20 in dPBS with 2% milk and added to the ELISA plate followed by 1hour incubation at 37°C. Plates are then washed six times followed by incubation with anti-human IgG HRP conjugated secondary antibody added at a 1:3000 dilution. Plates are washed a last time then incubated with developer followed by measuring the OD-450 value by our plate-reader. To determine antibody concentration, a linear relationship based on the human IgG standard was created in excel to determine the exact human ng/ml concentration within our supernatant and animal serum.

### Mouse infection with MA SARS-CoV-2 CMA4 strain

6-week-old female BALB/C mice were I.M. injected with mRNA-LNPs which encoded for the HC and LC of LY1404 or 76E1, respectively, or with eLNP as a negative control. HC and LC mRNA-LNPs were injected at 1:1 molar ratio with a combined mRNA dose of 15µg per animal. 24 hours after mRNA administration, all mice were infected intranasally with a 1 × 10^4^ PFU dose of MA SARS-CoV-2 CMA4 strain. Infected mice were monitored daily for morbidity. 2 days post viral infection, all mice were euthanized, and the mouse lungs were collected for the quantification of virus titers by plaque assay as detailed below.

### Plaque assay for MA-SARS-CoV-2 CMA4 infectious titers in lung

Vero E6 cells were seeded in 6-well plates and incubated at 37°C. Lung tissue homogenates were serially diluted (10-fold) in DMEM with 2% FBS and 0.2 ml was used to infect cells at 37°C for 1 hour. After the incubation, samples were overlaid with 2X-MEM (Gibco) with 8% FBS and 1.6% agarose (Promega). After 48 hour, plates were stained with 0.05% neutral red (Sigma-Aldrich) and plaques were counted to calculate virus titers expressed as PFU/ml.

### Hamster challenge with SARS-CoV-2 Omicron variant BQ.1

6-week-old male Syrian gold hamsters (HsdHan:Aura; Inotiv) were intramuscularly injected with mRNA-LNPs that respectively encoded the HC and LC chain of LY1404 or 76E1, or with eLNP as a negative control. HC and LC mRNA-LNPs were injected at 1:1 molar ratio with a combined RNA dose of 15µg per animal. At 24 hours post mRNA administration, all hamsters were challenged intranasally with a 2x10^4^ pfu/ml infectious dose of SARS-CoV-2 Omicron BQ.1. Animals were monitored daily for morbidity. At 2 days after viral infection, hamsters were euthanized, and lungs were collected for measuring viral titers and for histological analysis. The NW was collected prior to euthanizing as a means to measure viral titers as well in the URT.

### Quantification of BQ.1 viral titers in hamster airways by plaque assay

Homogenized lung tissue and collected NW was initially spiked and diluted out to 10^6^ in maintenance media. Diluted samples are then dispensed onto a confluent monolayer of Vero E6 cells (ATCC; CRL-1586) in a 96-well plate for 1 hour at 37°C with 5% CO_2_. After one hour, a complete overlay is applied to the cell surface and allowed to incubate for 48 hours, after which 10% formalin is added to the plate for fixation. Plates were then removed from BSL-3 to BSL-2 and monolayers washed with dPBS (Sigma) and incubated in permeabilization buffer consisting of dPBS supplemented with 0.1% bovine serum albumin (Sigma-Aldrich) and 0.1% saponin (Sigma-Aldrich) for 30 minutes at room temperature. Permeabilization buffer was removed, and monolayers were coated with rabbit polyclonal antibody against SARS-CoV N protein and incubated at 4°C overnight (S. Makino, Department of Microbiology and Immunology, UTMB) diluted in permeabilization buffer (1:3000). Excess antibody was washed away and monolayers were incubated for 1 hour at room temperature with HRP-conjugated goat anti-rabbit IgG (Cell Signaling Technology, 7040) diluted in permeabilization buffer (1:2000). Plates were washed one last time and foci were stained using KPL TrueBlue Peroxidase Substrate (SeraCare). Once foci were visible under a light microscope, water was dumped on to the plate for rapidly stopping the reaction. Plates were then read using a microplate reader and foci counted

### Lung histopathology

Lungs harvested from hamsters were fixed in 10% formalin and embedded in paraffin, sectioned and stained with hematoxylin and eosin Y using a protocol as previously reported ^(34)^. The slides were evaluated by a pathologist in a blinded manner.

### Statistics

Statistical analysis was performed using GraphPad Prisim 10 software. A one-way ANOVA was used for comparing the differences between the three groups (LY1404, 76E1 and eLNP) and for generating reported P-values. P-values <.05 were deemed significant.

## ACKNOWLEDGEMENT

We thank P. A. Valdes at UTMB for assistance in acquiring and preparing histopathological images. The research was in part supported by NIH grant AI181134. H.H. as investigator was supported by NIH grants AI157852 and AI181134. N.C.H. was supported by a NIH T32 training grant AI060549.

